# Identification of Southeast Asian *Anopheles* mosquito species with matrix-assisted laser desorption/ionization time-of-flight mass spectrometry using a cross-correlation approach

**DOI:** 10.1101/2024.07.10.602996

**Authors:** Victor Chaumeau, Sunisa Sawasdichai, Thu Zar Ma Ma Moe Min, Thithiwarada Kularbkeeree, Naw Jaruwan, Naw Gloria, Naw Yu Lee, Muesuwa Trackoolchengkaew, Monticha Phanaphadungtham, Patcharamai Rongthong, Aritsara Inta, Wanitda Watthanaworawit, François Nosten

## Abstract

Matrix-assisted laser desorption/ionization time-of-flight mass spectrometry (MALDI-TOF MS) is proposed for mosquito species identification. The absence of public repositories for sharing mass spectra and of open-source data analysis pipelines for fingerprint matching to mosquito species limits widespread use of this technology. The objective of this study was to develop an open-source data analysis pipeline for *Anopheles* species identification with MALDI-TOF MS. Malaria mosquitos were captured in 33 villages in Karen (Kayin) state in Myanmar. 359 specimens were identified with DNA barcodes and assigned to 21 sensu stricto species and 5 sibling species pairs or complexes. 3584 mass spectra of the head of these specimens identified with DNA barcoding were acquired and the similarity between mass spectra was quantified using a cross-correlation approach adapted from the published literature. A simulation experiment was carried out to evaluate the performance of species identification with MALDI-TOF MS at varying thresholds of cross-correlation index for the algorithm to output an identification result and with varying numbers of technical replicates for the tested specimens, considering PCR identification results as the reference. With one spot and a threshold value of −14 for the cross-correlation index on the log scale, the sensitivity was 0.99 (95%CrI: 0.98 to 1.00), the predictive positive value was 0.99 (95%CrI: 0.98 to 0.99) and the accuracy was 0.98 (95%CrI: 0.97 to 0.99). It was not possible to directly estimate the sensitivity and negative predictive value because there was no true negative in the assessment. In conclusion, the modified cross-correlation approach can be used for matching mass spectral fingerprints to predefined taxa and MALDI-TOF MS is a valuable tool for rapid, accurate and affordable identification of malaria mosquitos.

## Introduction

The Thailand-Myanmar border is a rural area of Southeast Asia where people and animals living on both sides of the border are exposed to diseases transmitted by anopheline mosquitos (Mullen and Durden, 2009; Rattanarithikul et al., 2006). Due to specificities in the border ecology, the diversity of mosquito species is among the highest worldwide (Morgan et al., 2013). Endemic *Anopheles* species are distributed among 18 main groups within the 3 subgenera *Anopheles*, *Baimaia* and *Cellia* (Rattanarithikul et al., 2006). The main vectors are *An. minimus* (Minimus Complex, Funestus Group), *An. maculatus*, *An. sawadwongporni* (Maculatus Group), *An. dirus* and *An. baimaii* (Dirus Complex, Leucosphyrus Group). *An. pseudowillmori* (Maculatus Group), *An. aconitus* (Aconitus Subgroup, Funestus Group) and some species of the Annularis and Barbirostris Complexes are secondary vectors. Many other species are suspected vectors because they transmit malaria elsewhere but their vector status is not well characterized in this region (Chaumeau et al., 2018; Sinka et al., 2011). Accurate vector identification is important to assess transmission dynamics and evaluate vector-control interventions. However, closely related species can have overlapping morphological characters or be completely isomorphic thereby delimiting cryptic or sibling species complexes that cannot be distinguished by morphology (Manguin et al., 2008).

Matrix-assisted laser desorption/ionization time-of-flight mass spectrometry (MALDI-TOF MS) is used for the identification of living organisms including medically important pathogens and arthropods (Calderaro and Chezzi, 2024; Croxatto et al., 2012). Several studies have striven to develop reference mass spectra databases and data analysis methods for *Anopheles* species identification with MALDI-TOF MS (Briolant et al., 2020; Diarra et al., 2019; Nabet et al., 2021; Raharimalala et al., 2017; Tandina et al., 2018), including Southeast Asian species (Chaumeau et al., 2024; Huynh et al., 2022; Mewara et al., 2018). However, the lack of open-source data analysis pipeline and sharing of raw mass spectra limits widespread use of this technology. Several methods were proposed to match mass spectral fingerprints of biological specimens to predefined taxa. Signature peaks can be used to distinguish between taxa but only little information is retained in the analysis and this approach is not reliable with complex matrix and if numerous closely related taxa are to be identified. Alternatively, fingerprint matching can be performed by measuring similarity between mass spectra. Predictive modeling such as linear discriminant analysis or random forest can identify groups of more similar mass spectra corresponding to specimens of the same species (Loaiza et al., 2019; Müller et al., 2013; Satten et al., 2004), but overfitting and the context-specific nature of the similarity metrics they produce can be problematic. Another method is to compute pairwise similarity scores between pairs of mass spectra and to select the spectra pair that has the highest similarity in a query of a reference mass spectra database with a panel of test samples to be identified. This can be implemented with the MALDI Biotyper® (Bruker Daltonics, Bremen, Germany) and Vitek® system (bioMérieux, Marcy L’Etoile, France) commercial software (Calderaro and Chezzi, 2024), or with the more recently developed free MSI2 application available online (Chaumeau et al., 2024), but the matching algorithms were not disclosed. Arnold and Reilly have proposed a mathematical algorithm to quantify similarity between MALDI-TO MS mass spectra with a cross-correlation approach and used it to distinguish between 25 strains of *Escherichia coli* (Arnold and Reilly, 1998), but there has been no assessment of this method to identify mosquito species.

The objective of this study was to develop an open-source analysis pipeline for the identification of *Anopheles* species of the Thailand-Myanmar border with MALDI-TOF MS, allowing rapid, accurate and affordable identification of mosquitos collected during entomological surveys. *Anopheles* mosquito specimens collected in Karen (Kayin) state in Myanmar were identified with DNA barcodes and subsequently analyzed with MALDI-TOF MS. A reference mass spectra database was constructed using open-source software and a cross-correlation algorithm based on the seminal work of Arnold and Reilly (Arnold and Reilly, 1998). The performance of identification with MALDI-TOF MS was evaluated using an external dataset generated from specimens captured in the same area. Eventually, the database was upgraded to include all available mass spectra (whether they were initially included in the reference or in the test panel). All analysis code and data are available online, allowing anyone to replicate the methodology and create a reference mass spectra database in-house.

## Material and methods

### Samples collection and preparation

Entomological surveys were carried out in 33 villages in Karen (Kayin) state in Myanmar between November 9^th^, 2020 and October 10^th^, 2022. Mosquitos were captured into 5-mL plastic tubes using the human-landing catch and animal-baited trap (buffalo, cow or goat) collection methods. Upon reception in the laboratory (1 to 10 days after collection), mosquitos were sorted macroscopically at the genus level and *Anopheles* specimens were identified at the group level using a dichotomic morphological identification key (Rattanarithikul et al., 2006). A subset of these malaria mosquitos was randomly selected to constitute a panel representative of the different villages and species diversity. Specimens assigned to the reference panel were dissected on the reception day. The dissected head was stored at −80°C in 1.5 mL microcentrifuge tubes for 85 to 422 days before being analyzed with MALDI-TOF MS. The remaining anatomical parts were taken into 1.5-mL microcentrifuge tubes containing silica beads and stored at −20°C for a similar time before being analyzed with PCR. Specimens assigned to the test panel were kept into 1.5-mL microcentrifuge tubes containing silica beads and stored at −20°C for 392 to 1069 days before being dissected and analyzed with PCR and MALDI-TOF MS.

### DNA extraction and PCR amplification

DNA was extracted from dissected mosquito abdomens using the cetyl trimethylammonium bromide method as described previously (Chaumeau et al., 2016). Amplification of cytochrome c oxidase subunit I (COI) was performed using the primer pair LCO1490 (5’-GGT CAA CAA ATC ATA AAG ATA TTG G-3’) and MTRN (5’-AAA AAT TTT AAT TCC AGT TGG AAC AGC-3’) (Folmer et al., 1994; Kumar et al., 2007). The PCR mix was composed of 1X Goldstar™ DNA polymerase (Eurogentec, Seraing, Belgium, catalog number: PK-0064-02) and 400 nM of each primer. The PCR was conducted in a total reaction volume of 25 μl (4 μl of DNA template diluted at the 1/100^th^ in PCR grade water and 21 μl of PCR mix). The thermocycling protocol consisted in an initial activation step of 1 min at 94 °C, followed by 40 amplification cycles of 20 sec at 94 °C, 20 sec at 51 °C and 30 sec at 72 °C. Reactions that failed to amplify the target were repeated with the reverse primer HCO2198 (5’-TAA ACT TCA GGG TGA CCA AAA AAT CA-3’) (Folmer et al., 1994) using the same reaction conditions. Amplification of internal transcribed spacer 2 (ITS2) was performed using the primer pair ITS2A (5’-TGT GAA CTG CAG GAC ACA T-3’) and ITS2B (5’-ATG CTT AAA TTY AGG GGG T-3’) (Beebe and Saul, 1995) with the same reaction conditions, except primer concentration which was 100 nM each. PCR products were purified using the illustra™ ExoProStar 1-Step commercial kit (cytiva, Marlborough, USA, catalog number: US77720) following manufacturer’s instructions. Sanger sequencing of the purified product was outsourced to Macrogen™ (Seoul, South Korea) and performed with the forward primer (accession numbers: PP339876-PP340064, PP372871-PP373055, PP968979-PP969128 and PP976491-PP976529). Samples that could not be sequenced were removed from the panel.

### Protein extraction and mass spectra acquisition

Dissected heads were put into 1.5-mL microcentrifuge tubes and rinsed in 70% ethanol for 10 min. The tubes were centrifuged at 18,000 g for 10 min and the supernatant was discarded. After a second centrifugation at 18,000 g for 2 min, the remaining ethanol solution was removed using a micropipette and left to evaporate until dry. Protein extraction was performed by adding 10 μL of 70% formic acid solution (bioMérieux, Lyon, France, catalog number: 411072). After manual homogenization with a micropipette, the homogenates were incubated for 5 min at room temperature. Then, 10 μL of 100% acetonitrile (VWR, Randor, USA, catalog number: 20060.32) were added and the samples were incubated for an additional 5 min. The homogenates were then centrifuged at 18,000 g for 2 min, and 1 μL of the protein extracts was deposited onto a disposable target plate (bioMérieux, catalog number: 410893). Once dried, the deposits were covered with 1 μL of alpha-cyano4-hydroxycinnamic acid matrix (bioMérieux, catalog number: 411071). Ten spots were made for each specimen. Mass spectra were acquired with a Vitek® system (bioMérieux) in the research use only mode using the Shimadzu Biotech Launchpad MALDI-TOF MS application (Shimadzu Biotech, Kyoto, Japan). The spectra were acquired in linear mode in the ion-positive mode at a laser frequency of 60 Hz and a mass range of 2000-20000 Da. *Escherichia coli* ATCC 8739 was used as a control calibration spot for each run following manufacturer’s instruction. Raw mass spectra were exported in mzXML file format and used for subsequent analysis.

### *Anopheles* species identification with DNA sequence data

Sanger chromatograms were manually trimmed and inspected using Unipro UGENE software version 48 (Okonechnikov et al., 2012) to trim the beginning and the end of each sequence and correct artifactual polymorphisms. ITS2 was annotated using a 5.8S-28S rRNA interaction and hidden Markov model-based annotation program available online (Keller et al., 2009). COI and ITS2 sequences were queried against the NCBI Nucleotide collection (nr/nt) database using BLASTn (Altschul et al., 1990) and matched to a mosquito species when the identity between the query and subject sequences was ≥ 98%. COI sequences were also queried against the Barcode of Life Data System (BLOD), which includes a larger collection of COI sequences than the NCBI Nucleotide database (Ratnasingham and Hebert, 2007). To further validate BLAST and BOLD identification results, a phylogenetic analysis was conducted using the study sequences and reference sequences sourced from Genbank. COI sequences were aligned with Clustal W version 2.1 (Larkin et al., 2007) and ITS2 sequences were aligned with MAFFT using the X-INS-i algorithm and default parameters (Katoh and Toh, 2008). The phylogenetic analysis was performed in MrBayes v3.2, using a general time-reversible substitution model and gamma rates (Ronquist et al., 2012). In the analysis of COI sequences, the dataset was partitioned to estimate different mutation rates for the two first and the third codon positions. Each analysis comprised two independent runs with four chains running for 1,000,000 generations with a sample frequency of 100 generations. The first 25% trees were discarded as burn-in, and posterior probabilities were estimated from the remaining trees to infer branch support.

### Construction of the reference mass spectra database

Spectra were visualized and processed using R software version 4.2 and the MALDIquant package (Gibb and Strimmer, 2012; R Core Team, 2013). Processing of raw mass spectra included intensity transformation and smoothing, baseline removal, normalization of intensity values and spectra alignment (Figure 1). A cross-correlation index (CCI) was then calculated for each pairwise spectra comparison using a custom algorithm adapted from the work of Arnold and Reilly (Arnold and Reilly, 1998). The algorithm outputs the maximum value of the cross-correlation function for two input mass spectra over specified mass intervals, and the CCI is given by the product of the local cross-correlation value of each mass interval. If no local maximum is detected on a given mass interval, the algorithm returns 0. Therefore, the range of possible CCI values on the log scale (log_10_[CCI]) is a real number varying between a negative infinite limit (too dissimilar spectra) and 0 (identical spectra). In this assessment, the CCI algorithm was parameterized with mass intervals of 500 Da spanning the mass range 3000-12000 Da.

**Figure 1.**
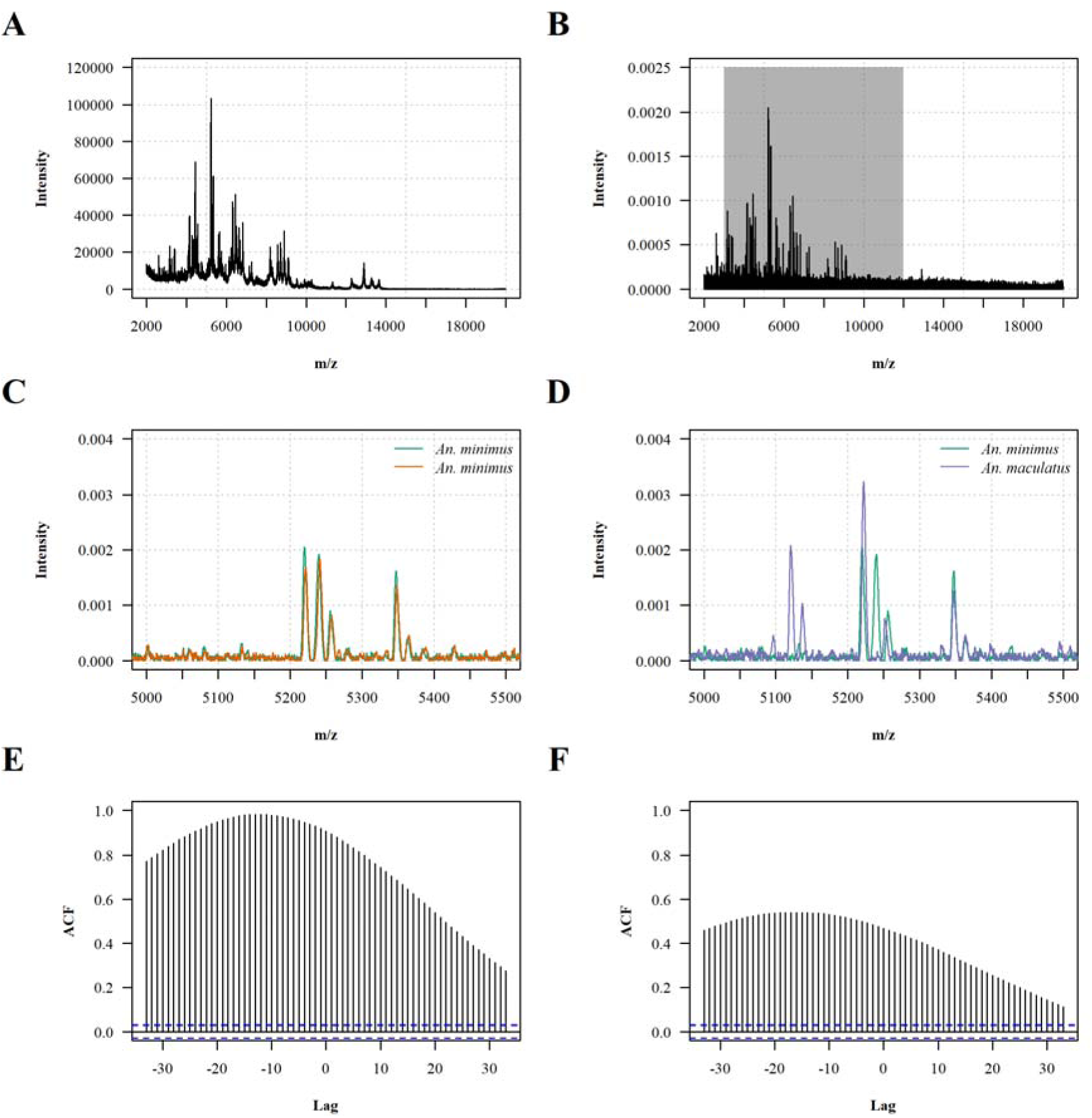
Principle of the cross-correlation algorithm. (A) A typical raw MALDI-TOF mass spectrum of *An. minimus*. (B) The corresponding processed spectrum after intensity normalization and baseline removal. (C) Comparison of the processed mass spectra of two *An. minimus* specimens over the 5000-5500 kDa mass interval showing high similarity between spectra. (D) Comparison of the processed mass spectra of an *An. minimus* specimen and of an *An. maculatus* specimen over the 5000-5500 kDa mass interval showing limited similarity between spectra. (E) The cross-correlation function of the two *An. minimus* spectra over the 5000-5500 kDa mass interval gives a local maximum of 0.982. (F) The cross-correlation function of the *An. minimus* and *An. maculatus* spectra over the 5000-5500 kDa mass interval gives a local maximum of 0.540. If no local maximum of the cross-correlation function is detected, the algorithm is parameterized to return 0. The resulting cross-correlation index on the log scale (log_10_[CCI]) over the 3000-12000 kDa mass range is −4.9 for the two *An. minimus* spectra and -Infinite for the *An. minimus* and *An. maculatus* spectra.

### Evaluation of the performance of MALDI-TOF MS identification

In order to evaluate the performance of *Anopheles* species identification with MALDI-TOF MS, the test panel was queried against the reference mass spectra database. A simulation experiment was carried out by selecting at random a specified number of spots per specimen of the test panel varying from 1 to 9 with 1000 repeats. The MALDI-TOF identification result was defined as the reference spectrum giving the highest CCI value, considering all randomly selected spots in the analysis. An identification threshold was set for the CCI value below which no identification result was given. The MALDI-TOF identification result was categorized as true positive (test specimen matching with the same species in the reference mass spectra database with a CCI value above the identification threshold), true negative (specimen of a species not represented in the reference mass spectra database with a CCI value below the identification threshold), false positive (matches between different species with a CCI value above the identification threshold) and false negative (species represented in the reference mass spectra database with a CCI value below the identification threshold), considering PCR identification results as the reference. The results of this simulation experiment were used to estimate the sensitivity, positive predictive value and accuracy. Noteworthy, as species coverage in the reference mass spectra database increases with the sample size, very few specimens of species not referenced in the reference mass spectra database are included in the test panel thereby impairing the estimation of specificity and negative predictive value (because there is no true negative result in the assessment). Therefore, a separate analysis was carried out after excluding comparisons of the same species (i.e., forcing the result to be either falsely positive or truly negative by mimicking queries of unreferenced species), allowing indirect estimation of the specificity but only for queries of unreferenced species (i.e., the probability of a result being falsely positive if the query species is not referenced in the reference database) but not of the negative predictive value (because the negative predictive value also depends on the number of false negative).

### Ethics

The study was approved by the Oxford Tropical Research Ethics Committee, the Karen Department of Health and Welfare, Karen National Union and the Tak Province Border Community Ethics Advisory Board (Cheah et al., 2010).

## Results

### Panel composition

403 *Anopheles* specimens were selected for inclusion in either the reference or the test panel (270 and 133 specimens, respectively). 254/270 (94%) specimens of the reference panel could be identified based on the analysis of ITS2 (32 specimens), COI (24 specimens) or both markers (198 specimens). Given the limited added value of COI sequencing for the identification of *Anopheles* species of the reference panel, only ITS2 sequencing was carried out for the specimens of the test panel and 105/133 (79%) specimens could be identified. Based on these DNA sequence data, 359/403 (89%) specimens were assigned to 26 taxa including 21 sensu stricto species and 5 sibling species pairs or complexes, and were subsequently analyzed with MALDI-TOF MS (Table 1 and Figure 2). Two specimens identified as *An. baimaii*/*dirus* based on COI sequencing were not included in the above figures because other specimens of *An. dirus* and *An. baimaii* were identified based on ITS2 sequencing. Four specimens of the Barbirostris group for which only a portion of the ITS2 could be sequenced were also removed because the query cover in the BLAST analysis was too short for reliable species identification (approximately 50 base pairs). One specimen of *An. annularis* s.l. had a COI sequence with 97.9% identity with available *An. annularis* s.l. COI DNA barcodes (below the identification threshold of BOLD system) and branched separately from other *An. annularis* s.l. specimens in the phylogenetic tree. Therefore, it was identified as a putatively new sibling species in this complex based on this marker. However, in the absence of additional evidence to support this taxonomic assignment and given only one specimen was identified, this assignment was merged with *An. annularis* s.l. in subsequent analysis. Similarly, two clades of *An. minimus* were identified based on COI DNA sequences but no distinction was made between both clades.

**Figure 2.**
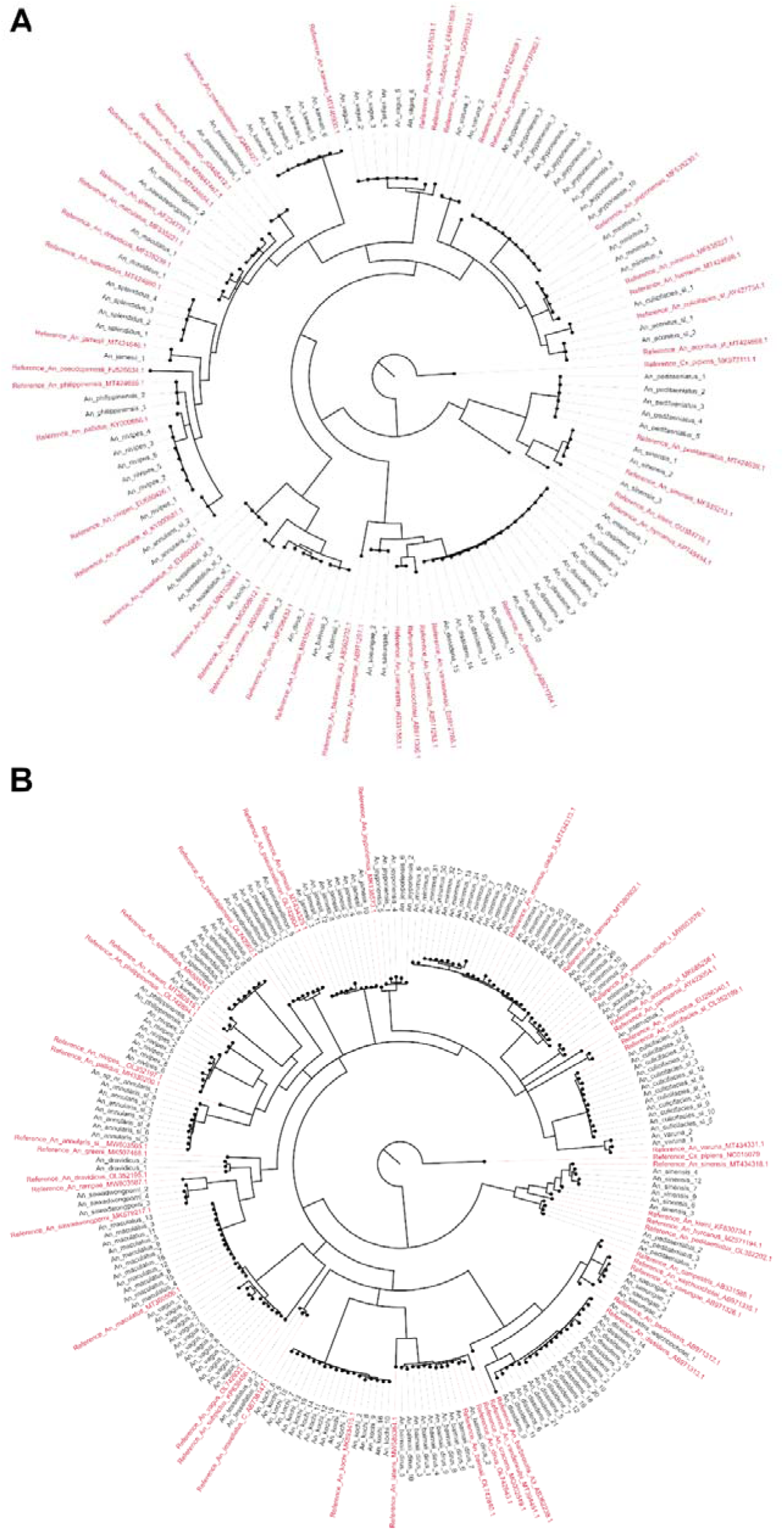
Phylogenetic trees based on Bayesian inference for the *Anopheles* specimens included in the panel. (A) Consensus tree for the ITS2 sequences. (B) Consensus tree for the COI sequences. Reference sequences sourced from GenBank are showed in red.

**Table 1.**
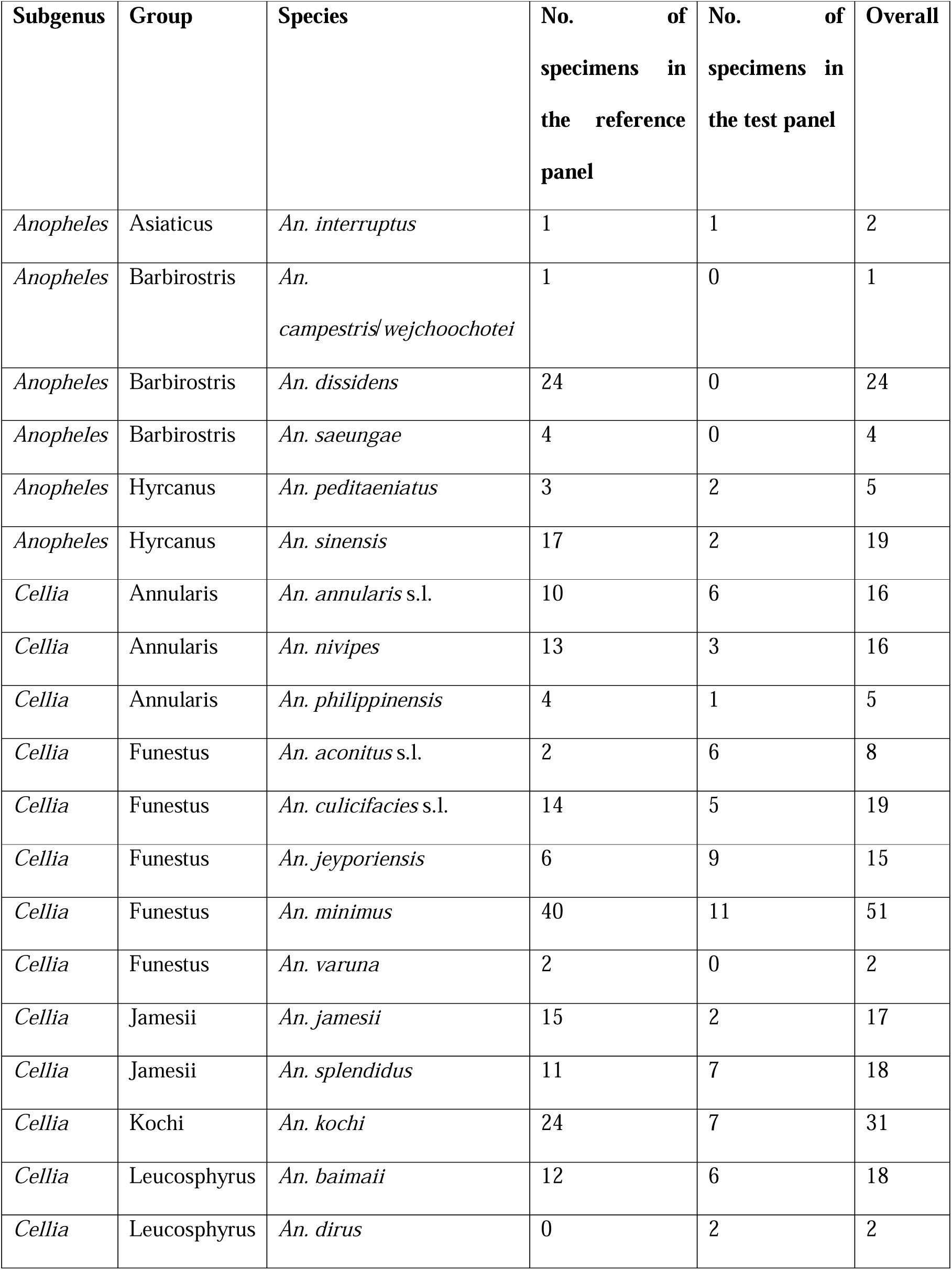
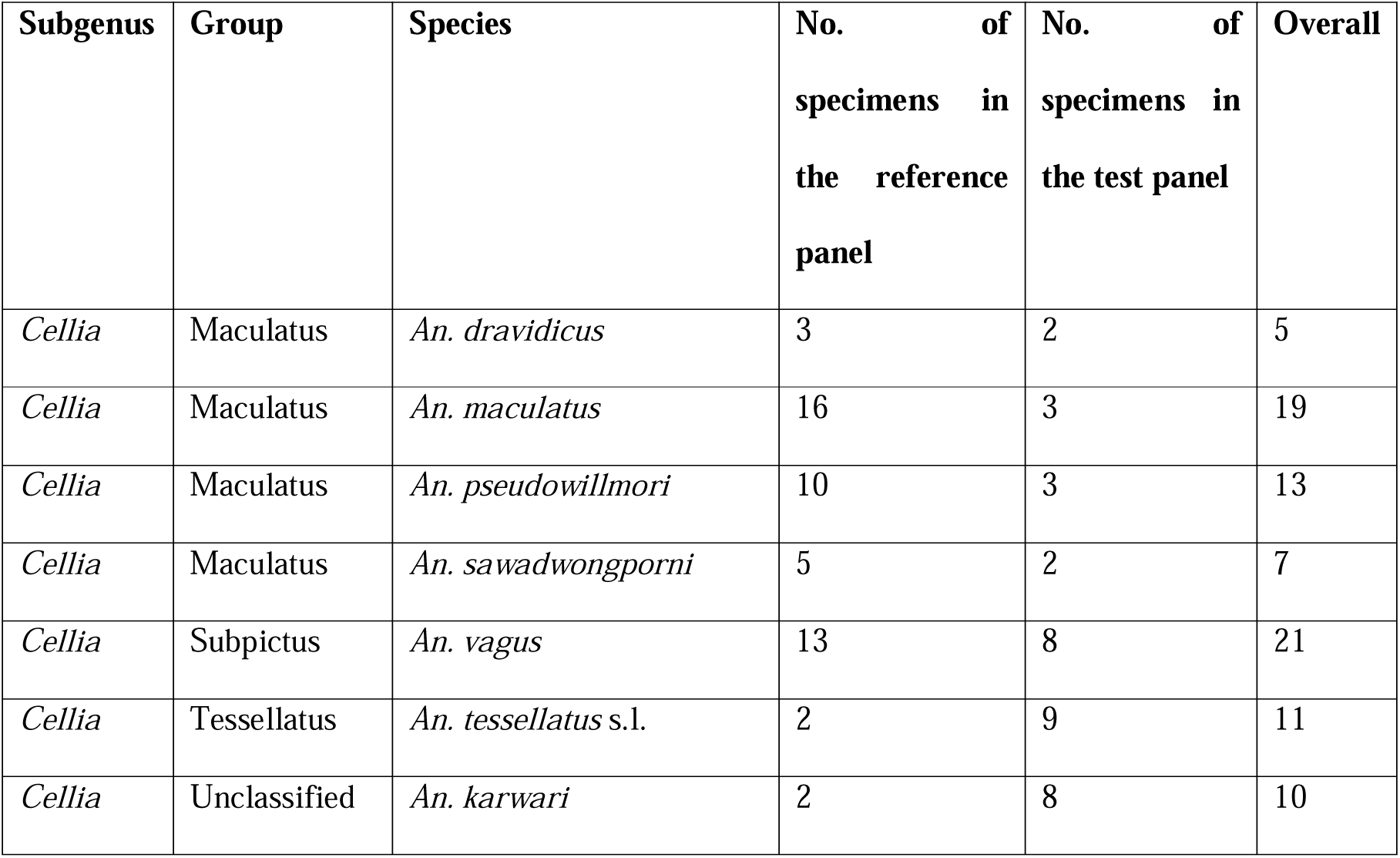
Panel composition.

| Subgenus | Group | Species | No. of specimens in the reference panel | No. of specimens in the test panel | Overall |
| --- | --- | --- | --- | --- | --- |
| *Anopheles* | Asiaticus | *An. interruptus* | 1 | 1 | 2 |
| *Anopheles* | Barbirostris | *An. campestris/wejchoochotei* | 1 | 0 | 1 |
| *Anopheles* | Barbirostris | *An. dissidens* | 24 | 0 | 24 |
| *Anopheles* | Barbirostris | *An. saeungae* | 4 | 0 | 4 |
| *Anopheles* | Hyrcanus | *An. peditaeniatus* | 3 | 2 | 5 |
| *Anopheles* | Hyrcanus | *An. sinensis* | 17 | 2 | 19 |
| *Cellia* | Annularis | *An. annularis* s.l. | 10 | 6 | 16 |
| *Cellia* | Annularis | *An. nivipes* | 13 | 3 | 16 |
| *Cellia* | Annularis | *An. philippinensis* | 4 | 1 | 5 |
| *Cellia* | Funestus | *An. aconitus* s.l. | 2 | 6 | 8 |
| *Cellia* | Funestus | *An. culicifacies* s.l. | 14 | 5 | 19 |
| *Cellia* | Funestus | *An. jeyporiensis* | 6 | 9 | 15 |
| *Cellia* | Funestus | *An. minimus* | 40 | 11 | 51 |
| *Cellia* | Funestus | *An. varuna* | 2 | 0 | 2 |
| *Cellia* | Jamesii | *An. jamesii* | 15 | 2 | 17 |
| *Cellia* | Jamesii | *An. splendidus* | 11 | 7 | 18 |
| *Cellia* | Kochi | *An. kochi* | 24 | 7 | 31 |
| *Cellia* | Leucosphyrus | *An. baimaii* | 12 | 6 | 18 |
| *Cellia* | Leucosphyrus | *An. dirus* | 0 | 2 | 2 |
| *Cellia* | Maculatus | *An. dravidicus* | 3 | 2 | 5 |
| *Cellia* | Maculatus | *An. maculatus* | 16 | 3 | 19 |
| *Cellia* | Maculatus | *An. pseudowillmori* | 10 | 3 | 13 |
| *Cellia* | Maculatus | *An. sawadwongporni* | 5 | 2 | 7 |
| *Cellia* | Subpictus | *An. vagus* | 13 | 8 | 21 |
| *Cellia* | Tessellatus | *An. tessellatus* s.l. | 2 | 9 | 11 |
| *Cellia* | Unclassified | *An. karwari* | 2 | 8 | 10 |

### Construction of the reference mass spectra database

2535 mass spectra of the 254 *Anopheles* specimens selected for inclusion in the reference panel and identified with PCR were acquired, yielding 3,211,845 pairwise comparisons of distinct spectra pairs. The reference mass spectra database covered 25 taxa including 20 sensu stricto species and 5 sibling species pairs or complexes. A CCI value reflecting similarity between spectra was computed for every spectra pairs. The repeatability, reproducibility and specificity of mass spectra was assessed using subsets of the comparisons (Figure 3): the median log_10_[CCI] was −7.9 (inter-quartile range [IQR]: −9.2 to −6.8) for comparisons of technical replicates of the same specimen, −10.7 (IQR: −12.6 to −9.4) for comparisons of different specimens of the same species and -Infinite (IQR: -Infinite to - Infinite) for comparisons of different species. Exclusion of spectra with low repeatability or reproducibility had no significant effect on the performance of MALDI-TOF MS identification (Appendix Table S1), therefore all spectra were kept in subsequent analysis. The heatmap grid of the median log_10_[CCI] collated by specimen included in the reference database, despite showing a high similarity between sibling species of the Barbirostris Complex and between some species of the Neomyzomyia Series, confirmed the high repeatability, reproducibility and specificity of mass spectra (Figure 4). Hierarchical clustering of mass spectra showed that specimens of the same species mostly grouped in the same branch but limited concordance with species phylogeny (Appendix Figure S1). When considering only the highest CCI value with self-matching disabled, 2477/2535 (97.7%) of the spectra included in the reference database matched with the same species (median log_10_[CCI]: −7.8 [IQR: −8.8 to −7.0]) and 58/2535 (2.3%) spectra matched with another species (Figure 5 and Appendix Table S2). Among the mismatches, 19 were spectra of species represented by only one specimen and thus not included in the queried dataset because self-matching was disabled (median log_10_[CCI]: −13.4 [IQR: −14.1 to −9.6]), and 39 were true cross-matches between two referenced species (median log_10_[CCI]: −11.7 [IQR: - 13.6 to −9.8]). Interestingly, the distribution of CCI values in concordant matches varied across taxa suggesting variable level of inter-specimen reproducibility and possible cryptic diversity in some taxa (Appendix Table S3). True cross-matches were observed between *An. dissidens* and *An. saeungae* (2 spectra, log_10_[CCI]: −9.1 and −7.0) and vice versa (5 spectra, median log_10_[CCI]: −7.0 [IQR: −7.5 to −7.0]), between *An. dravidicus* and *An. maculatus* (4 spectra, median log_10_[CCI]: −9.7 [IQR: −10.0 to −9.6]), *An. minimus* (1 spectrum, log_10_[CCI]: - 10.0), *An. nivipes* (1 spectrum, log_10_[CCI]: −10.2) and *An. splendidus* (1 spectrum, log_10_[CCI]: −9.9), between *An. karwari* and *An. dissidens* (1 spectrum, log_10_[CCI]: −11.8) and *An. jamesii* (1 spectrum, log_10_[CCI]: −9.9), between *An. kochi* and *An. pseudowillmori* (1 spectrum, log_10_[CCI]: −17.0), between *An. pseudowillmori* and *An. maculatus* (2 spectra, log_10_[CCI]: −11.7 and −11.9), between *An. sawadwongporni* and *An. maculatus* (8 spectra, median log_10_[CCI]: −13.8 [IQR: −14.0 to −13.6]) and *An. nivipes* (2 spectra, log_10_[CCI]: −14.5 and −13.9), and between *An. tessellatus* s.l. and *An. kochi* (10 spectra, median log_10_[CCI]: −12.2, IQR: −13.1 to −11.0). Among the species represented by only one specimen, 8/9 *An. campestris*/*wejchoochotei* spectra matched with *An. dissidens* (median log_10_[CCI]: −9.4 [IQR: −9.7 to −9.2]) and the remaining spectrum matched with *An. saeungae* (log_10_[CCI] = 10.2), and 10/10 *An. interruptus* spectra matched with *An. minimus* (median log_10_[CCI]: −14.1 [IQR: −14.5 to −13.8]).

**Figure 3.**
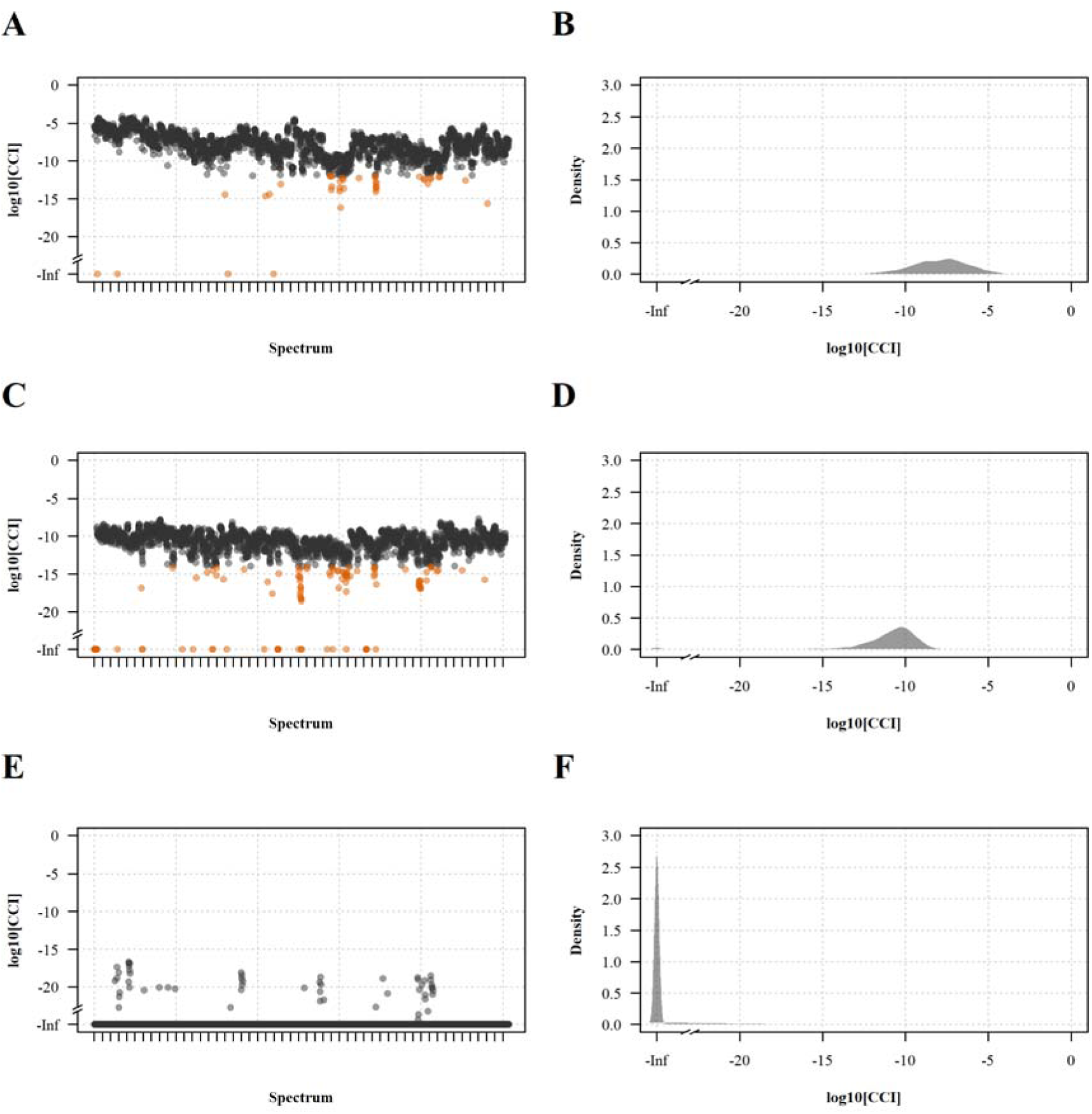
Repeatability, reproducibility and specificity of the mass spectra. (A) Median log_10_[CCI] of pairwise comparisons between technical replicates of the same specimen collated by mass spectrum; (B) the corresponding density function. (C) Median log_10_[CCI] of pairwise comparisons between spectra of different specimens of the same species collated by mass spectrum; (D) the corresponding density function. (E) Median log_10_[CCI] of pairwise comparisons between spectra of different species collated by mass spectrum; (F) corresponding density function.

**Figure 4.**
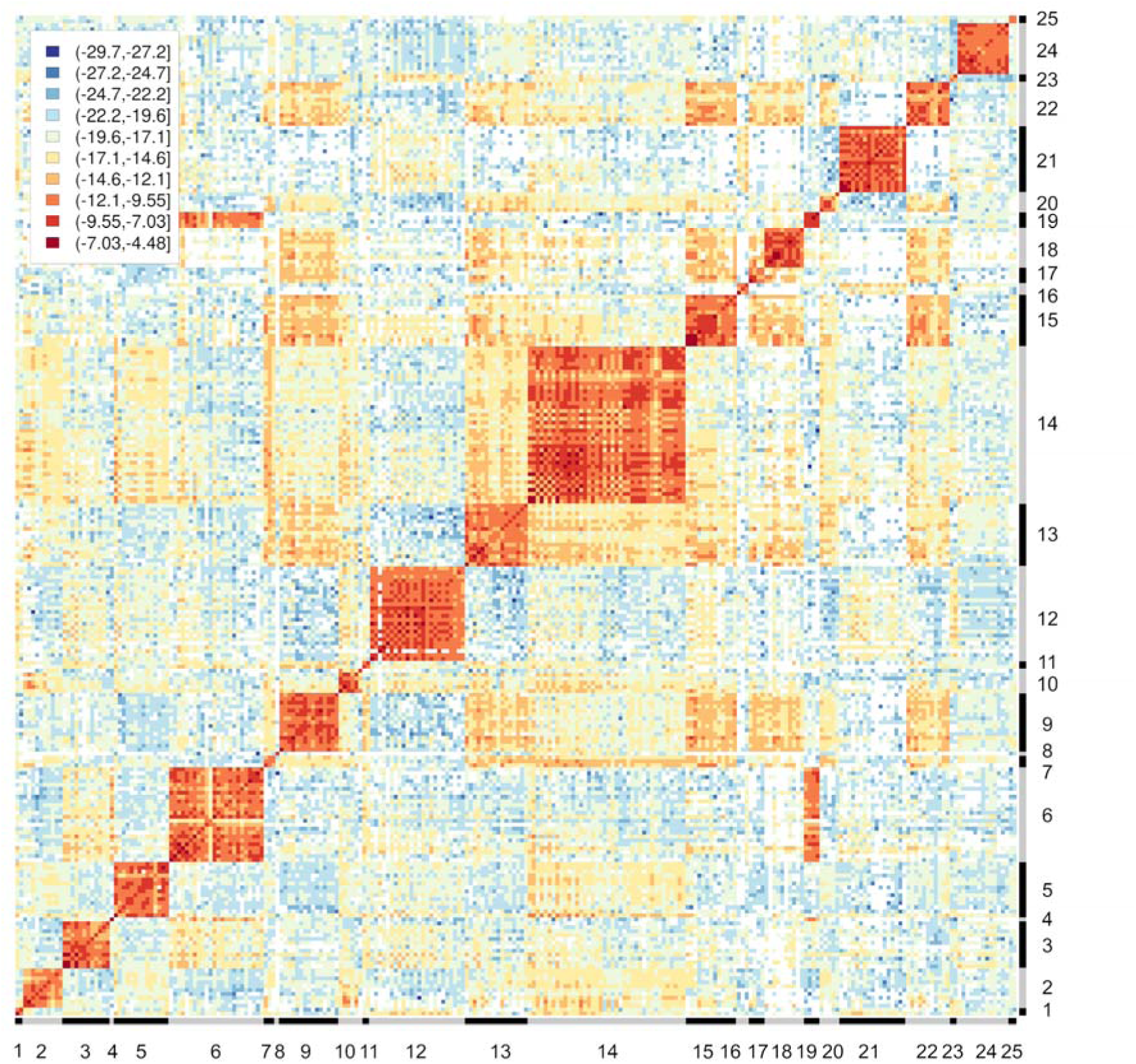
Heat map grid of the median cross-correlation index collated by specimen included in the reference mass spectra database. Red color on the central diagonal and the orange color around the central diagonal show the high repeatability and reproducibility of mass spectra, respectively. The blue color out of the central diagonal shows the high specificity of mass spectra. Orange color out of the central diagonal shows the high similarity between sibling species of the Barbirostris Complex and some species of the Neomyzomyia Series. Negative infinite values are showed in white. 1: *An. aconitus* s.l.; 2: *An. annularis* s.l.; 3: *An. baimaii*; 4: *An. campestris*/*wejchoochotei*; 5: *An. culicifacies* s.l.; 6: *An. dissidens*; 7: *An. dravidicus*; 8: *An. interruptus*; 9: *An. jamesii*; 10: *An. jeyporiensis*; 11: *An. karwari*; 12: *An. kochi*; 13: *An. maculatus*; 14: *An. minimus*; 15: *An. nivipes*; 16: *An. peditaeniatus*; 17: *An. philippinensis*; 18: *An. pseudowillmori*; 19: *An. saeungae*; 20: *An. sawadwongporni*; 21*: An. sinensis*; 22*: An. splendidus*; 23: *An. tessellatus* s.l.; 24: *An. vagus*; 25: *An. varuna*.

**Figure 5.**
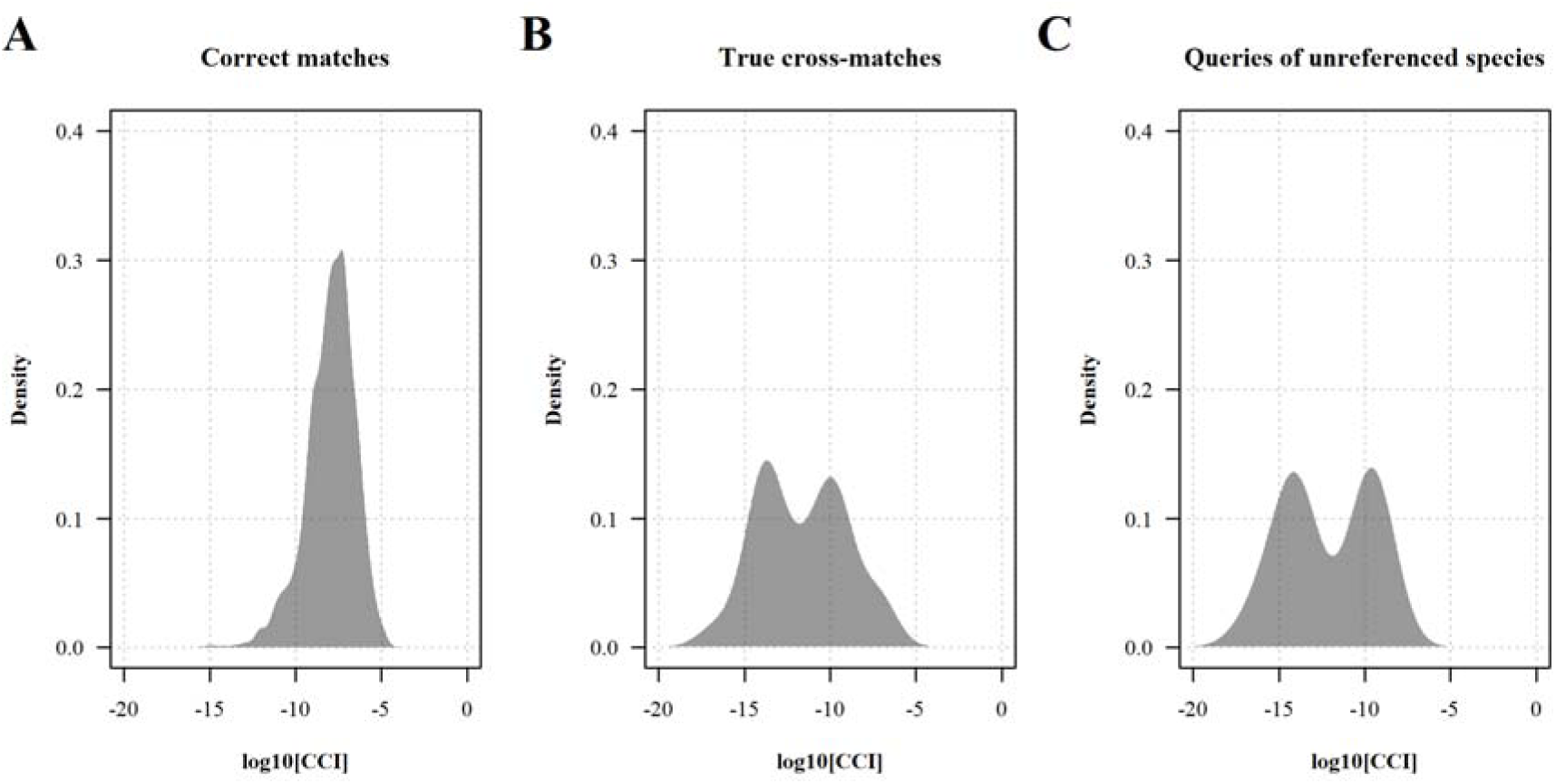
Distribution of the maximum log_10_[CCI] value collated by spectra in bank-to-bank comparison by category of result, excluding comparisons between technical replicates of the same sample. (A) Correct matches with another specimen of the same species. (B) True cross-matches between a species referenced in the queried database and a different species. (C) Species represented by only one specimen in the reference mass spectra database and therefore not included in the queried database because self-matching was disabled.

### Evaluation of the performance

In order to evaluate the performance of *Anopheles* species identification with MALDI-TOF MS, 1049 mass spectra of the 105 PCR-identified specimens included in the test panel were queried against the reference database, yielding 2,659,215 pairwise comparisons. The sensitivity, positive predictive value and accuracy were estimated at varying values of identification threshold and numbers of technical replicates by carrying out a simulation experiment with 1000 repeats. In this experiment, a subset of the spectra was selected at random for each specimen of the test panel and the identification result was defined as the reference mass spectra giving the highest CCI value considering all randomly selected replicates for a given test specimen. Noteworthy, true negative results become rare as species coverage increase in the reference database thus impairing the estimation of the specificity and negative predictive value. Therefore, the specificity was assessed by disabling matches between specimens of the same species thereby mimicking queries of unreferenced species only and forcing the result to be either truly negative of falsely positive (see Material and Methods). Increasing the number of technical replicates increased the sensitivity and decreased the specificity resulting in an increased classification accuracy, whereas increasing the identification threshold toward a more stringent value decreased the sensitivity and increased the specificity resulting in a decreased classification accuracy (Table 2 and Figure 6). Setting the identification threshold to −14 (low stringency) and considering one spot in the analysis, the sensitivity was 0.96 (95% credible interval [CrI]: 0.92 to 0.99), the specificity was 0.39 (95%CrI: 0.32 to 0.46), the positive predictive value was 0.94 (95%CrI: 0.92 to 0.96) and the accuracy was 0.90 (95%CrI: 0.87 to 0.94). With 4 spots, the sensitivity was 1.00 (95%CrI: 1.00 to 1.00), the specificity 0.19 (95%CrI: 0.16 to 0.23), the positive predictive value 0.93 (95%CrI: 0.91 to 0.95) and the accuracy 0.93 (95%CrI 0.92 to 0.93). When considering an identification threshold of −12 (high stringency) and either 1 or 4 spots in the analysis, the sensitivity was 0.82 (95%CrI: 0.77 to 0.87) versus 0.92 (95%CrI: 0.89 to 0.94), the specificity 0.85 (95%CrI: 0.80 to 0.90) versus 0.71 (95%CrI: 0.68 to 0.76), the positive predictive value was 0.97 (95%CrI: 0.94 to 0.99) versus 0.95 (95%CrI: 0.94 to 0.97) and the accuracy was 0.81 (95%CrI: 0.75 to 0.86) versus 0.88 (95%CrI: 0.85 to 0.9), respectively. The list of spectra that gave a false positive or false negative result considering an identification threshold of −14 and all simulation repeats is provided in the Appendix Table S4.

**Figure 6.**
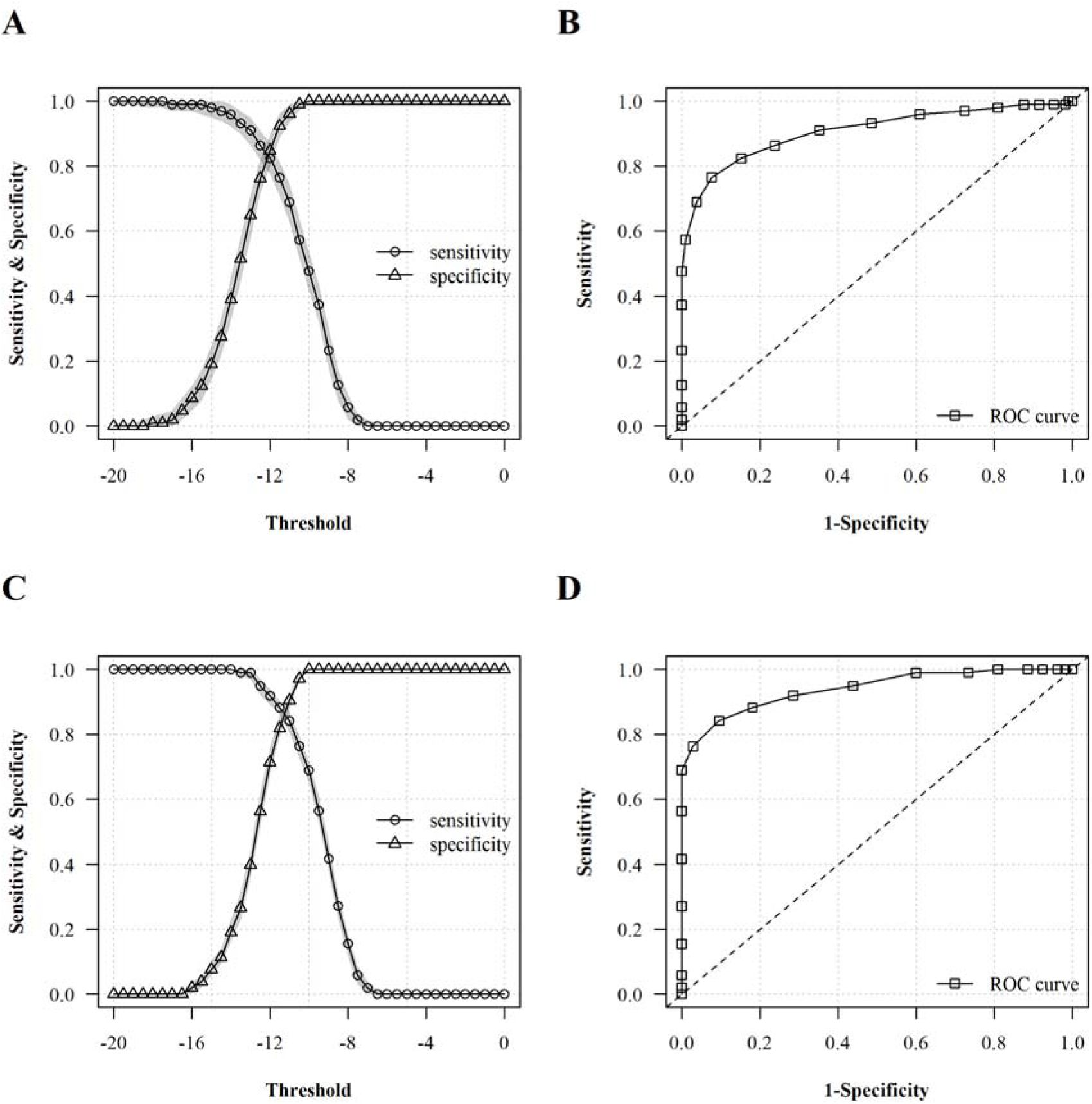
Evaluation of the performance of the reference mass spectra database for *Anopheles* species identification using the test panel. (A) Sensitivity and specificity estimated at varying identification threshold considering one spot per specimen; (B) corresponding receiving operator characteristics curve. (C) Sensitivity and specificity estimated at varying identification threshold considering four spots per specimen; (D) corresponding receiving operator characteristics curve. The shaded areas in panels A and C show the 95% credible interval around the median value of 1000 simulations. The dashed line in panels B and D shows the performance of a random classification.

**Table 2.** Performance of *Anopheles* species identification with MALDI-TOF MS.

| **Threshold** | **No.**<br>**spots** | **Sensitivity** | **Specificity** | **Positive predictive**<br>**value** | **Accuracy** |
| --- | --- | --- | --- | --- | --- |
| -14 | 1 | 0.96 (0.92 to 0.99) | 0.39 (0.32 to 0.46) | 0.94 (0.92 to 0.96) | 0.9 (0.87 to 0.94) |
| -14 | 2 | 1 (0.98 to 1) | 0.27 (0.22 to 0.31) | 0.94 (0.91 to 0.96) | 0.93 (0.9 to 0.96) |
| -14 | 4 | 1 (1 to 1) | 0.19 (0.16 to 0.23) | 0.93 (0.91 to 0.95) | 0.93 (0.91 to 0.95) |
| -14 | 9 | 1 (1 to 1) | 0.14 (0.14 to 0.16) | 0.92 (0.92 to 0.93) | 0.92 (0.92 to 0.93) |
| -13 | 1 | 0.91 (0.86 to 0.95) | 0.65 (0.58 to 0.71) | 0.95 (0.93 to 0.97) | 0.87 (0.83 to 0.9) |
| -13 | 2 | 0.96 (0.93 to 0.99) | 0.51 (0.46 to 0.58) | 0.95 (0.92 to 0.97) | 0.91 (0.88 to 0.94) |
| -13 | 4 | 0.99 (0.97 to 1) | 0.4 (0.36 to 0.45) | 0.93 (0.91 to 0.96) | 0.92 (0.9 to 0.95) |
| -13 | 9 | 1 (0.99 to 1) | 0.3 (0.3 to 0.32) | 0.92 (0.92 to 0.93) | 0.92 (0.91 to 0.93) |
| -12 | 1 | 0.82 (0.77 to 0.87) | 0.85 (0.8 to 0.9) | 0.97 (0.94 to 0.99) | 0.81 (0.75 to 0.86) |
| -12 | 2 | 0.89 (0.86 to 0.92) | 0.78 (0.73 to 0.83) | 0.96 (0.94 to 0.98) | 0.86 (0.83 to 0.89) |
| -12 | 4 | 0.92 (0.89 to 0.94) | 0.71 (0.68 to 0.76) | 0.95 (0.94 to 0.97) | 0.88 (0.85 to 0.9) |
| -12 | 9 | 0.95 (0.93 to 0.95) | 0.62 (0.61 to 0.65) | 0.94 (0.94 to 0.95) | 0.9 (0.88 to 0.9) |

### Database upgrade

Finally, the reference database was upgraded in order to include all PCR-identified specimens (whether they were initially included in the reference or in the test panel) and the same analysis was carried out. The upgraded database was composed of 3584 spectra of 359 specimens including 21 sensu stricto species and 5 sibling species pairs or complexes (Table 1). In bank-to-bank analysis with self-matching disabled, 3528/3584 (98.4%) of the spectra matched with the same species (median log_10_[CCI]: −8.3 [IQR: −9.5 to −7.3]) and 56/3584 (1.6%) with another species (47 were true cross-matches, median log_10_[CCI]: −12.4 [IQR: −13.8 to −10.1] and 9 were queries of species represented by only one specimen in the upgraded database, median log_10_[CCI]: −9.5 [IQR: −10.1 to −9.2], Appendix Table S5). Setting the identification threshold to −14 and considering 1 spot in the analysis, the sensitivity was 0.99 (95%CrI: 0.98 to 1.00), the specificity (for queries of unreferenced species) was 0.18 (95%CrI: 0.16 to 0.21), the predictive positive value was 0.99 (95%CrI: 0.98 to 0.99) and the accuracy was 0.98 (95%CrI: 0.97 to 0.99, Appendix Table S6).

## Discussion

A reference mass spectra database for the identification of *Anopheles* mosquito species of the Thailand-Myanmar border with MALDI-TOF MS was developed. The database references 21 sensu stricto species and 5 sibling species pairs or complexes, including the main malaria vectors in this area. Fingerprint matching of *Anopheles* species was carried out with a custom algorithm based on the seminal work of Arnold and Reilly who first described the principle of the cross-correlation index as a quantitative measure of similarity between mass spectra (Arnold and Reilly, 1998). In this assessment, the identification performances were excellent with a sensitivity of 0.99 (95%CrI: 0.98 to 1.00), a predictive positive value of 0.99 (95%CrI: 0.98 to 0.99) and an accuracy of 0.98 (95%CrI: 0.97 to 0.99) considering an identification threshold value for the log_10_[CCI] of −14 and only one technical replicate per test specimen to be identified. Noteworthy, queries of unreferenced species become rare as species coverage in the reference database increases (Roswell et al., 2021), thus impairing the estimation of specificity and negative predictive value (because there is no true negative in the assessment). By disabling matches between spectra of the same species (i.e., mimicking queries of unreferenced species and forcing the MALDI-TOF MS identification result to be either false positive or true negative), the estimated sensitivity was only 0.18 (95%CI: 0.16 to 0.21). This finding suggests that using an identification threshold with low stringency could give poor identification performance if the species composition of the test panel is different than that of the reference mass spectra library, and highlights the importance of exhaustive species referencing in the reference mass spectra database or concomitant assessment of species diversity in a subset of the samples to be identified with MALDI-TOF MS with a DNA sequencing approach.

In this setting, about a hundred mosquito specimens can be identified with MADLI-TOF MS per day for a cost of approximately USD 1.5 per sample. The main limiting factor of throughput are the manual method used to extract proteins and the relatively long acquisition time. Future studies may strive to address these limitations. The main cost driver was the use of disposable slides (30 % of the total cost) because there are no commercially available reusable slides for the MALDI-TOF MS device used in this study. This finding highlights the importance of considering the cost of consumables when procuring of MALDI-TOF MS device, as it may vary a lot between device manufacturers and country. *Anopheles* species identification with MALDI-TOF MS has a higher taxonomic resolution than routine morphology, which can give a reliable identification result at the group or subgroup level for 500 to 800 specimens per person per day with reasonably achievable expertise in entomology for being deployed in virtually any setting. Morphological identification at the species or sibling species complex level requires a level of expertise in mosquito taxonomy that can be difficult to achieve, very good quality specimens and the throughput is less than for simple classification at the group or subgroup level. MALDI-TOF MS proved more reliable than the PCR assays used in this study: 98% of the PCR-identified specimens were accurately identified with MALDI-TOF MS (considering PCR as the reference) whereas only 89% of the specimens initially selected for inclusion in the panel could be identified with PCR despite repeated testing. Although the performance of PCR assays may be better in other settings, especially with well-optimized high-throughput DNA sequencing platform as was recently developed for the identification of mosquito species across the entire *Anopheles* genus (Boddé et al., 2022), this result demonstrates that very good identification performance can be achieved relatively easily with MALDI-TOF MS technology in settings with limited expertise in molecular biology.

This study has several limitations. Species coverage was not exhaustive and some specimens could not be identified at the species level. Therefore, the ability of the CCI algorithm to differentiate between sibling species remains largely undetermined. This limitation could be alleviated by carrying out additional testing of the remaining DNA extracts and by increasing the sampling efforts to reference additional species in the database. The performance of the database for the identification of specimens of the same species collected in different areas was not assessed. Additional mosquito collection and acquisition of reference mass spectra (i.e., high quality mass spectra from molecularly identified specimen) would be required to alleviate this limitation. The performance of MALDI-TOF MS identification with other anatomical parts was not assessed. This question was out of the scope of this study as it would have required a lot of additional testing. The effect of storage conditions and duration on the performance was not assessed. This would have required a different design that was also out of the scope of the current study.

Future research should be carried out to improve the code and develop an identification service readily available to end-users with limited bioinformatics skills, as was previously proposed with the MSI2 application (Chaumeau et al., 2024). Moreover, the performance and added value of *Anopheles* species identification with MALDI-TOF MS should be further validated at scale and head-to-head comparisons with other identification methods would be valuable. Non-malaria mosquitos include important vector species. Little is known about their ecology and biology in relation to disease transmission because of the challenges associated with sampling and identification (Chen et al., 2017; Maquart et al., 2021; Tangena et al., 2017; Thongsripong et al., 2013). Therefore, additional studies should be carried out to develop a reference mass spectra library for the identification of non-malaria mosquito species, and more generally of arthropod vectors of medical and veterinary importance, to support investigations of the diseases they transmit.

Accurate, fast and affordable vector species identification is paramount to investigate the dynamics of mosquito-borne disease transmission and evaluate the impact of control interventions. Despite decades of research, MALDI-TOF MS is seldom used operationally. This is largely due to the lack of open-source data analysis pipeline and data sharing. This study is an important step forward toward widespread use of MALDI-TOF MS for identification of arthropod vectors in resource-limited setting. In conclusion, MALDI-TOF MS is a promising tool for malaria vector identification but further research is needed to validate it at scale.

## Data availability statement

All analysis code and data are available via an accompanying github repository: https://github.com/victorSMRU/malditof-cci. The dataset was deposited on Zenodo: https://zenodo.org/records/12704052.

## Acknowledgements

We thank the staff of the Entomology, Laboratory, Malaria and Data Management Departments of the Shoklo Malaria Research Unit for their help with collection, processing and management of the samples and data included in this study. The Shoklo Malaria Research Unit is part of the Mahidol-Oxford Tropical Medicine Research Unit, supported by Wellcome, U.K. This research was funded by in part by Wellcome (#220211) and in part by the Bill and Melinda Gates Foundation (#OPP1177406). For the purpose of Open Access, the authors have applied a CC BY public copyright licence to any Author Accepted Manuscript version arising from this submission.

