## Supplementary figures and images for "Identification of Southeast Asian *Anopheles* mosquito species with matrix-assisted laser desorption/ionization time-of-flight mass spectrometry using a cross-correlation approach"

### figureS1_dendrogram.png

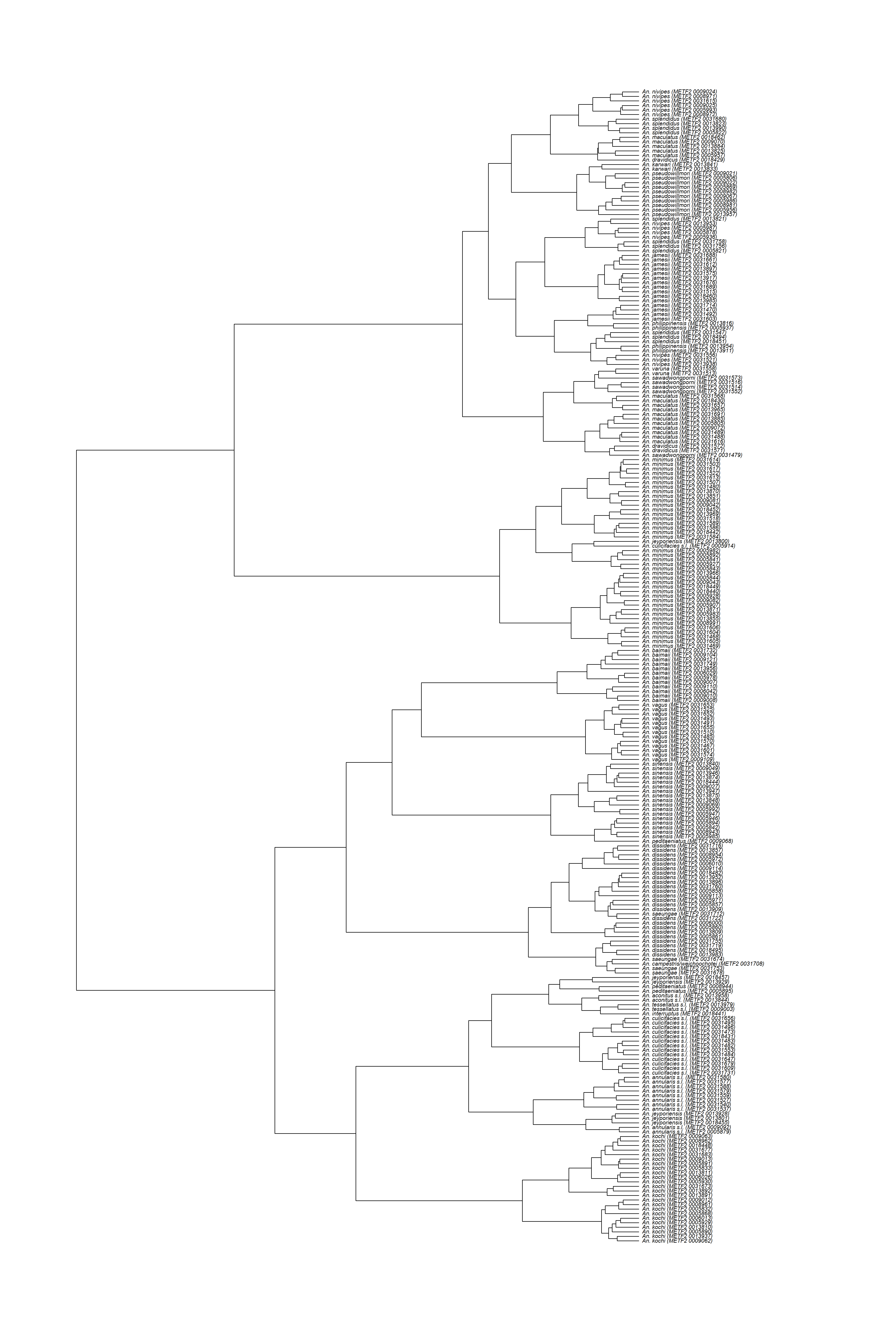
